# Phylogenetic signal is associated with the degree of variation in root-to-tip distances

**DOI:** 10.1101/2020.01.28.923805

**Authors:** Mezzalina Vankan, Simon Y.W. Ho, Carolina Pardo-Diaz, David A. Duchêne

**Affiliations:** Research School of Biology, Australian National University, ACT 2601, Australia; School of Life and Environmental Sciences, University of Sydney, Camperdown, NSW 2006, Australia; Biology Program, Universidad del Rosario, Carrera 24 No.63C-69, Bogotá, 111221 Colombia

**Keywords:** Phylogenomics, substitution rate, nucleotide substitution model, branch support, data filtering

## Abstract

The phylogenetic information contained in sequence data is partly determined by the overall rate of nucleotide substitution in the genomic region in question. However, phylogenetic signal is affected by various other factors, such as heterogeneity in substitution rates across lineages. These factors might be able to predict the phylogenetic accuracy of any given gene in a data set. We examined the association between the accuracy of phylogenetic inference across genes and several characteristics of branch lengths in phylogenomic data. In a large number of published data sets, we found that the accuracy of phylogenetic inference from genes was consistently associated with their mean statistical branch support and variation in their gene tree root-to-tip distances, but not with tree length and stemminess. Therefore, a signal of constant evolutionary rates across lineages appears to be beneficial for phylogenetic inference. Identifying the causes of variation in root-to-tip lengths in gene trees also offers a potential way forward to increase congruence in the signal across genes and improve estimates of species trees from phylogenomic data sets.

---

The phylogenetic signal in a molecular sequence alignment is influenced by a number of factors, including the substitution rate at which the sequences have evolved relative to the timescale of the process. In principle, the amount of information in the sequence alignment depends on the overall substitution rate of the gene (Goldman 1998; Xia et al. 2003; Townsend and Leuenberger 2011; Klopfstein et al. 2017; Steel and Leuenberger 2017). However, the substitution rate might be a poor predictor of the accuracy of the inferred tree topology (Aguileta et al. 2008). This is because the phylogenetic signal in a gene can be obscured by various forms of heterogeneity, such as variation in rates across sites (Su and Townsend 2015; Dornburg et al. 2019). Substantial rate heterogeneity can also be found across branches (Bromham and Penny 2003), but there is a still a limited understanding of the association between this form of rate variation and the topological signal in phylogenomic data sets.

Substitution rates can vary across genes and across lineages because of differences in selective pressures or limits on mutation rates (Gillespie 1991; Gaut et al. 2011). The factors that drive rate variation across genes and lineages can interact in what are known as “residual effects” (Gillespie 1991), potentially creating complex patterns of substitution rates across genes (Ho 2014; Duchêne and Ho 2015). Genes can also differ in their evolutionary histories, including their coalescence times, due to recombination breaking the linkage between sections of the genome (Maddison 1997). In addition to varying in their signals of rates and times, estimates of substitution rates in individual genes can be misled by a number of methodological factors, including model misspecification (Sullivan and Joyce 2005) and errors in alignment, orthology assignment, or sequencing (Wilkinson 1996; Sanderson and Shaffer 2002).

Any differences in evolutionary rates across genes will be reflected in the estimates of gene tree branch lengths. In statistical phylogenetic inference, branch lengths are closely linked to the estimate of tree topology. For example, long branches can have negative impacts on phylogenetic accuracy because of their tendency to be grouped together (“long-branch attraction”; Anderson & Swofford 2004). Even a single long branch can drastically change the phylogenetic signal in the data (Su and Townsend 2015). Meanwhile, low substitution rates can lead to a lack of phylogenetic information and even to a greater amount of phylogenetic error than in sequences that have evolved with very high substitution rates (Yang 1998). Although most research has focused on the differences in overall substitution rates across genes, the variation in the signal of rates across lineages is likely to provide a more nuanced and accurate predictor of the topological signal across the genome.

One potential predictor of phylogenetic accuracy is the degree of variation in the inferred distances from the root to each of the tips in a given gene tree. If substitution rates have been constant across lineages, the root-to-tip distances are expected to be proportional to time. Therefore, root-to-tip distances should all be identical in a data set where the samples come from the present and the sequences have evolved under a strict molecular clock. In theory, it is unlikely that any poor estimation in branch lengths will produce identical root-to-tip distances. Variation in root-to-tip distances might be caused by variation in rates across lineages, but critically, it is also diagnostic of the presence of factors causing inaccurate estimates of branch lengths.

Variation in root-to-tip distances will not be informative in cases where low information content is due to fast diversification events (over short time-periods) or where multiple lineages have changed in evolutionary rate simultaneously (an “epoch” model of rate variation). An alternative predictor of phylogenetic accuracy is the ratio of the lengths of internal branches to terminal branches, also known as stemminess (Fiala and Sokal 1985). Low stemminess is typically associated with a poor topological signal (e.g., Penny et al. 2001; Duchêne et al. 2018c), yet it is frequently observed in phylogenetic trees (e.g., Phillimore & Price 2008). Some explanations for low stemminess include rapid diversification events (McPeek 2008), sparse taxon sampling (Penny et al. 2001; Cusimano and Renner 2010), underparameterization of the substitution model (Revell et al. 2005), and deep gene coalescences relative to species divergence times (Maddison 1997; Degnan and Rosenberg 2009).

Testing the link between characteristics of branch lengths and estimates of tree topology across genes has potential benefits for the design of phylogenomic studies. One approach to carrying out a phylogenomic study is to employ a criterion to select genes for analysis, a practice known as “data filtering” or “gene shopping” (Molloy and Warnow 2018). Some of the criteria that have previously been used for data filtering include phylogenetic branch supports (Blom et al. 2016), the amount of missing data (Molloy and Warnow 2018), measures of substitution model adequacy (Duchêne et al. 2018c; Richards et al. 2018), and base composition (Dávalos and Perkins 2008; Martijn et al. 2018). It not clear which of these criteria is the most effective (Molloy and Warnow 2018), but it is likely that no single criterion is universally applicable (Reddy et al. 2017). Nonetheless, branch lengths provide an estimate of the amount of genetic change that is captured in a data set, so it is reasonable to surmise that they present a general predictor of the accuracy of estimates of tree topology (Klopfstein et al. 2017).

In this study, we explore the association between three branch-length metrics and estimates of tree topology across a collection of 34 phylogenomic data sets. When examining individual data sets, we find that the tree length is not the best predictor of phylogenetic information content among genes. Across the 34 data sets, we observe an association between the performance of phylogenetic inference and the variation in root-to-tip distances.

Phylogenomic studies are likely to benefit from considering the heterogeneity in rates across lineages for describing the signal of tree topology across loci.

## Materials and Methods

We collected a set of 34 phylogenomic data sets covering a wide range of taxa and data types (Table 1), including intron and exon regions, ultraconserved elements, and anchor-enriched regions. The original studies varied widely in their treatment of these data sets. For instance, some studies considered the trees from each of the codon positions of protein-coding genes independently. We followed the data treatments used in the original studies so that our analyses would reflect the approaches that have been used in practice.

**Table 1.**
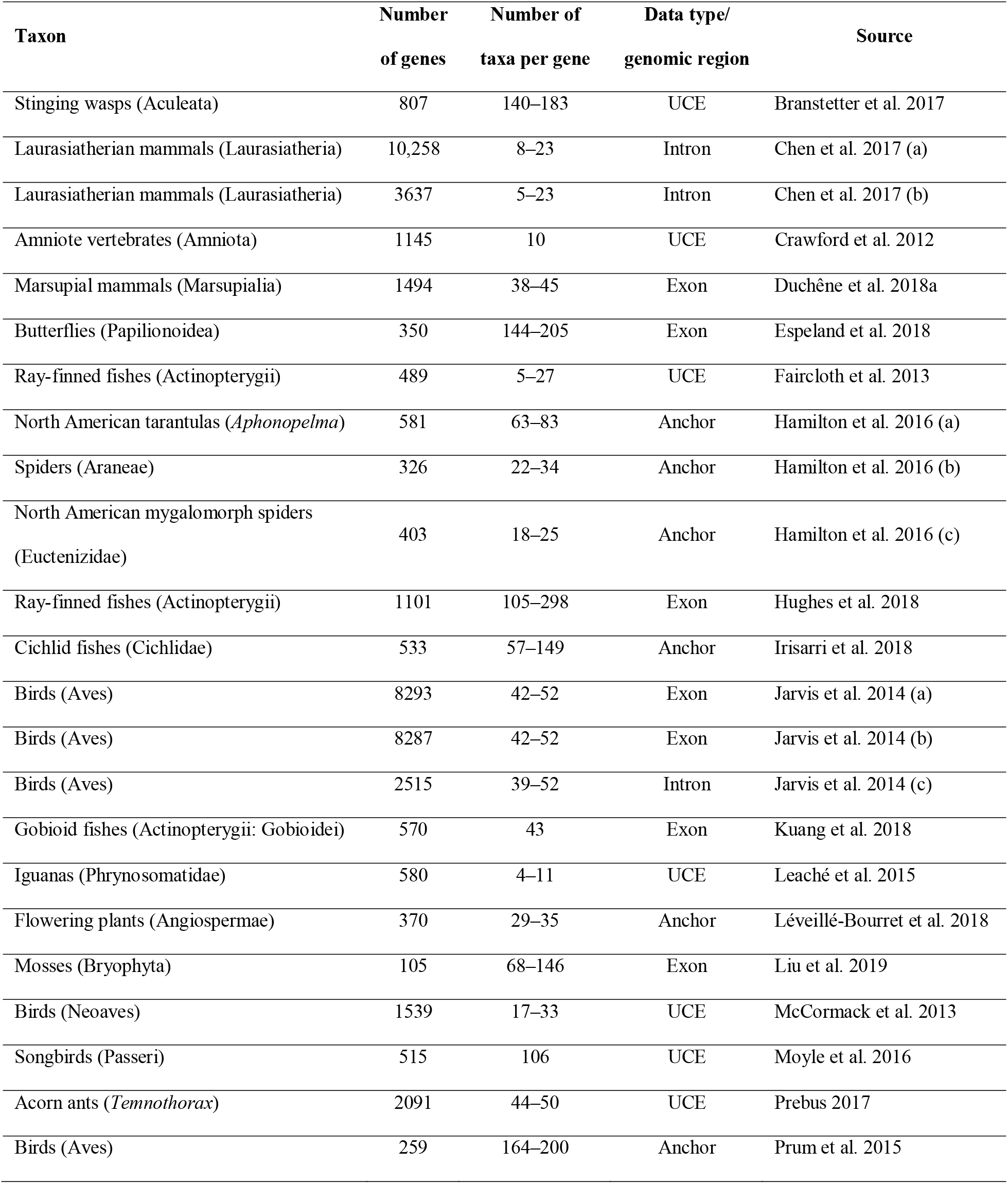
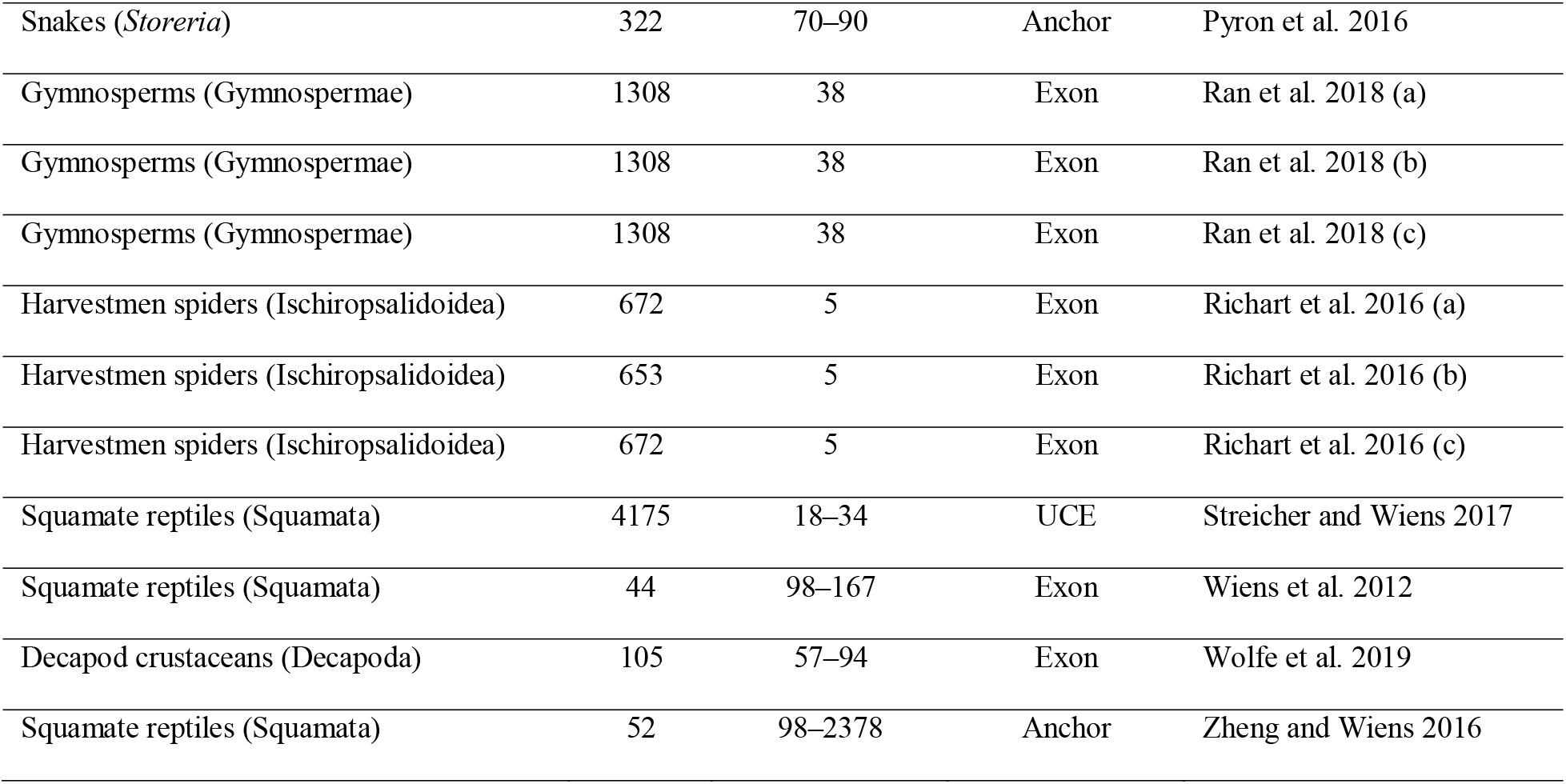
Phylogenomic data sets for which the association between phylogenetic signal and branch characteristics was tested. The treatment of data sets was similar to that in the original studies. Some of the published alignments were excluded because of numerical problems in phylogenetics software, excessive missing data, or file-format difficulties (such as those caused by unusual characters).

| Taxon | Number<br>of genes | Number of<br>taxa per gene | Data type/<br>genomic region | Source |
| --- | --- | --- | --- | --- |
| Stinging wasps (Aculeata) | 807 | 140–183 | UCE | Branstetter et al. 2017 |
| Laurasiatherian mammals (Laurasiatheria) | 10,258 | 8–23 | Intron | Chen et al. 2017 (a) |
| Laurasiatherian mammals (Laurasiatheria) | 3637 | 5–23 | Intron | Chen et al. 2017 (b) |
| Amniote vertebrates (Amniota) | 1145 | 10 | UCE | Crawford et al. 2012 |
| Marsupial mammals (Marsupialia) | 1494 | 38–45 | Exon | Duchêne et al. 2018a |
| Butterflies (Papilionoidea) | 350 | 144–205 | Exon | Espeland et al. 2018 |
| Ray-finned fishes (Actinopterygii) | 489 | 5–27 | UCE | Faircloth et al. 2013 |
| North American tarantulas (*Aphonopelma*) | 581 | 63–83 | Anchor | Hamilton et al. 2016 (a) |
| Spiders (Araneae) | 326 | 22–34 | Anchor | Hamilton et al. 2016 (b) |
| North American mygalomorph spiders<br>(Euctenizidae) | 403 | 18–25 | Anchor | Hamilton et al. 2016 (c) |
| Ray-finned fishes (Actinopterygii) | 1101 | 105–298 | Exon | Hughes et al. 2018 |
| Cichlid fishes (Cichlidae) | 533 | 57–149 | Anchor | Irisarri et al. 2018 |
| Birds (Aves) | 8293 | 42–52 | Exon | Jarvis et al. 2014 (a) |
| Birds (Aves) | 8287 | 42–52 | Exon | Jarvis et al. 2014 (b) |
| Birds (Aves) | 2515 | 39–52 | Intron | Jarvis et al. 2014 (c) |
| Gobioid fishes (Actinopterygii: Gobioidei) | 570 | 43 | Exon | Kuang et al. 2018 |
| Iguanas (Phrynosomatidae) | 580 | 4–11 | UCE | Leaché et al. 2015 |
| Flowering plants (Angiospermae) | 370 | 29–35 | Anchor | Léveillé-Bourret et al. 2018 |
| Mosses (Bryophyta) | 105 | 68–146 | Exon | Liu et al. 2019 |
| Birds (Neoaves) | 1539 | 17–33 | UCE | McCormack et al. 2013 |
| Songbirds (Passeri) | 515 | 106 | UCE | Moyle et al. 2016 |
| Acorn ants (*Temnothorax*) | 2091 | 44–50 | UCE | Prebus 2017 |
| Birds (Aves) | 259 | 164–200 | Anchor | Prum et al. 2015 |
| Snakes (*Storeria*) | 322 | 70–90 | Anchor | Pyron et al. 2016 |
| Gymnosperms (Gymnospermae) | 1308 | 38 | Exon | Ran et al. 2018 (a) |
| Gymnosperms (Gymnospermae) | 1308 | 38 | Exon | Ran et al. 2018 (b) |
| Gymnosperms (Gymnospermae) | 1308 | 38 | Exon | Ran et al. 2018 (c) |
| Harvestmen spiders (Ischiropsalidoidea) | 672 | 5 | Exon | Richart et al. 2016 (a) |
| Harvestmen spiders (Ischiropsalidoidea) | 653 | 5 | Exon | Richart et al. 2016 (b) |
| Harvestmen spiders (Ischiropsalidoidea) | 672 | 5 | Exon | Richart et al. 2016 (c) |
| Squamate reptiles (Squamata) | 4175 | 18–34 | UCE | Streicher and Wiens 2017 |
| Squamate reptiles (Squamata) | 44 | 98–167 | Exon | Wiens et al. 2012 |
| Decapod crustaceans (Decapoda) | 105 | 57–94 | Exon | Wolfe et al. 2019 |
| Squamate reptiles (Squamata) | 52 | 98–2378 | Anchor | Zheng and Wiens 2016 |

For each data set, we inferred the phylogeny using IQ-Tree (Nguyen et al. 2015) with the best-fitting substitution model from the GTR+Γ family. We then identified a set of gene trees from each data set that contained the same set of taxa. The taxon set was selected to maximize the product of the number of taxa and the number of genes, while maintaining full occupancy of the data matrix (for details see github.com/duchene/branch_length_influence_topology).

We calculated three test statistics that described the branch-length signal in each gene tree. These statistics included: (i) the coefficient of variation (CoV) in distances from the midpoint-root to the tips, which provides a measure of rate heterogeneity across lineages; (ii) tree length calculated as the sum of all branch lengths; and (iii) tree stemminess (Fiala and Sokal 1985). In addition, we calculated for each gene the mean of the statistical support across branches, using the Shimodaira-Hasegawa-like approximate likelihood-ratio test (aLRT; described in Anisimova and Gascuel 2006).

We assessed whether the four branch statistics could explain two different measures of the accuracy of tree topology estimates. The first measure was the topological distance from the species tree as estimated using a multispecies coalescent analysis in ASTRAL-III (Zhang et al. 2018) of the complete set of genes for the corresponding study. This evaluates the concordance between the phylogenetic signal in each gene tree and the underlying species history. The second measure of accuracy was the mean topological distance between the gene tree and all other gene trees from the corresponding data set. This evaluates the concordance of the signal in each gene tree with the remainder of the phylogenetic signals in the genome. All topological distances were calculated using the Robinson-Foulds topological distance (Robinson and Foulds 1981; Penny and Hendy 1985).

We used multiple linear regression to test whether the two measures of topological accuracy are explained by the four branch statistics. For each of the two response variables (topological distance of the gene tree to the species tree and mean topological distance to other gene trees), we first tested a model that included the complete data set of the genes from across the 34 studies (*N* = 36,075). We included the four branch statistics as explanatory variables in the regression models.

Since we aimed to identify the correlates of phylogenetic signal within each study, we attempted to account for the differences across studies in their results and their sample size. We included a random factor in each regression model that indicated the source study of each gene, this way accounting for the differences in patterns that might occur among studies. In this large model, we corrected tree length for the number of taxa by dividing it by the number of branches in the study (leading to the mean of branch lengths) to make the values fall on a similar scale across studies. We also explored the model when weighting each gene by the number of taxa in its source study, such that studies with a greater number of genes have a greater contribution to the model.

To focus further on the results within studies, we performed a second set of regression models where each study was examined independently. For each study, we tested whether our two response variables were explained by our four branch statistics. Therefore, this second set of analyses included two regression tests for each of the 34 studies that we examined. Tree length was left uncorrected for the number of branches in the regression models for individual studies.

## Results

The regression analyses that included the 34 complete data sets showed that some of our explanatory variables had a significant association with both measures of topological accuracy (topological distance to the species tree and topological distance to other gene trees; Fig. 1). Specifically, we found that topological accuracy has a positive association with the CoV in root-to-tip distances, and a negative association with mean aLRT branch support (Fig. 1). Mean aLRT branch support had the strongest association with both topological distance to the species tree and to other gene trees. Strikingly, we find limited evidence for an association between topological accuracy and tree length or stemminess. Results were comparable across regression models in which samples (genes) were weighted by number of branches or by number of taxa in respective studies (Supplementary Fig. S1).

**Figure 1.**
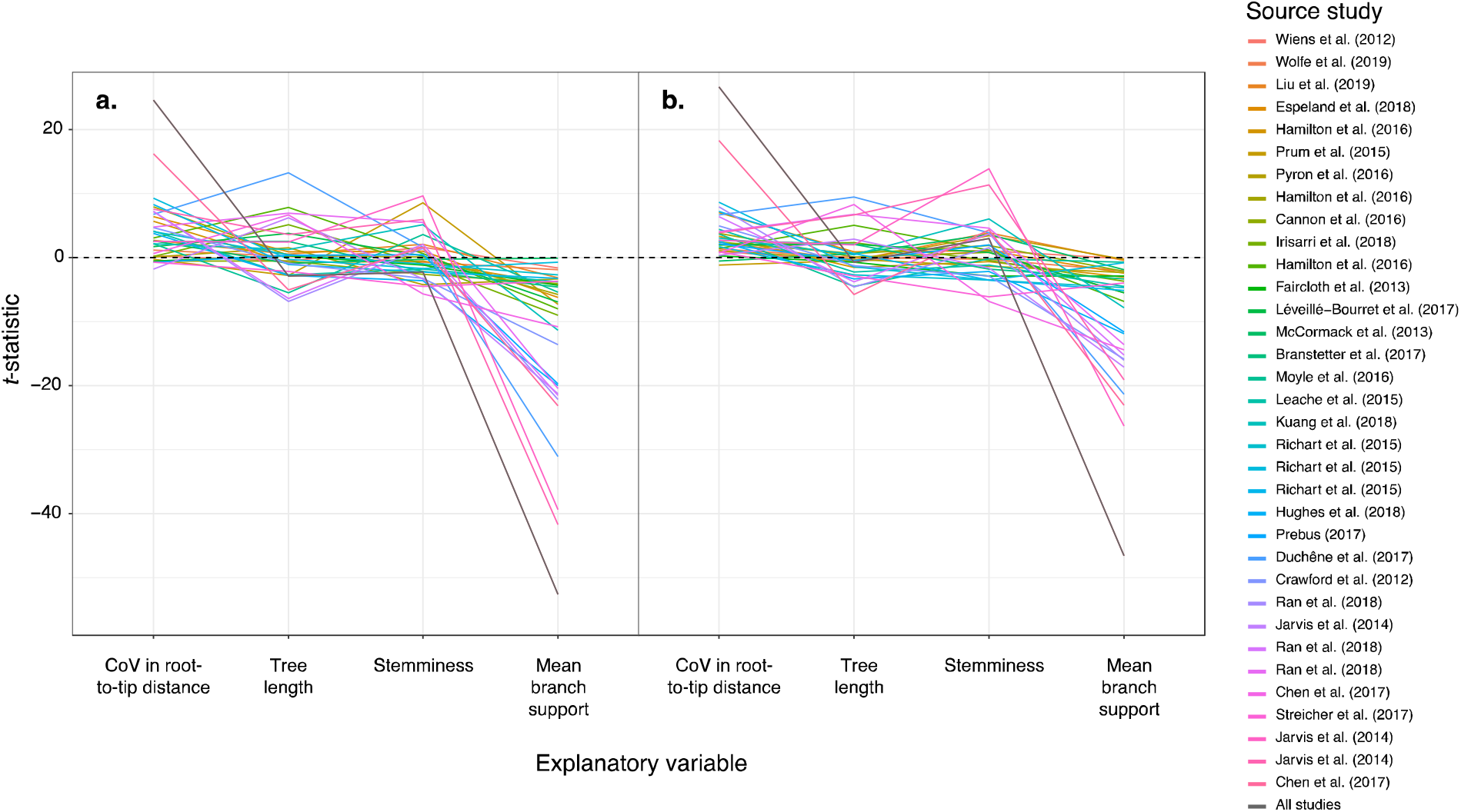
Summary *t*-statistic for multiple regression tests of the association between five explanatory variables describing branch lengths and each of two response variables: (a) topological distance between gene trees and the inferred species tree; and (b) mean distance from each gene tree to all other gene trees. The legend lists the source studies in ascending order of number of genes in the data set (see Table 1 for details).

The regression models that explored individual data sets supported the results from our larger regression models. Only a small minority of data sets showed an effect opposite to those observed for the CoV in root-to-tip distances and branch support. Meanwhile, there was substantial variation in terms of the association between topological accuracy and tree length or stemminess. As expected, the results of individual regression analyses showed greater *t*-statistics (smaller *P*-values) for data sets with large numbers of genes than for data sets with few genes. The *t*-statistics were comparable among regression models with each of the two measures of topological accuracy (Supplementary Fig. S2).

## Discussion

Our analyses of a collection of phylogenomic data sets have shown that low variation in root-to-tip distances and strong branch support in gene trees have a strong association with phylogenetic accuracy. Strikingly, tree length is a poor predictor of the accuracy of topological inference across gene trees. This is surprising because tree length is proportional to the overall substitution rate in a gene (Yang 1998), and is a prominent form of variation in the phylogenetic information across gene trees (Duchêne et al. 2020). These results are consistent with recent work that emphasized the importance of heterogeneity in the data rather than the overall substitution rate as an indicator of phylogenetic accuracy (Su and Townsend 2015; Dornburg et al. 2019).

Phylogenomic analysis can potentially be improved by focusing analyses and interpretation of results according to loci with particular patterns of rate variation across lineages. A formal method of identifying genes with constant rates across lineages is to compare a model of rate constancy versus one allowing rate variation (Felsenstein 1981). However, not all forms of rate variation across lineages are problematic for phylogenetics. One approach that might benefit phylogenomic studies is to identify the loci that have extreme patterns of rate variation among lineages and exclude them from analyses. Loci can then be retained for analysis when they contain patterns of rate variation across lineages that are mild and recurrent across multiple regions in the genome. Methods of describing the diversity of patterns of rate variation can be useful for this purpose (Duchêne et al. 2014).

Some of the extreme forms of variation in root-to-tip distances that lead to poor phylogenetic accuracy might be unrelated to variation in evolutionary rates across lineages. For example, sequence evolution might be heterogeneous across the tree, with variation in base composition or in transition probabilities among nucleotides (e.g., Dávalos & Perkins 2008; Foster *et al*. 2009; Martijn *et al*. 2018). Therefore, methods of assessing model adequacy are likely to be useful complementary diagnostics for improving the accuracy of topological inferences (Brown and ElDabaje 2009; Doyle et al. 2015; Höhna et al. 2017; Duchêne et al. 2018b, 2018c).

Variation in root-to-tip distances might also be an artefact of data preparation, rather than model performance. If model performance was a primary driver of phylogenetic accuracy, then we expect poor accuracy to be strongly associated with low stemminess (Revell et al. 2005). One wide-ranging solution to errors in data preparation is to identify and remove any taxa that have a highly variable position in a each given gene tree, also known as “rogue taxa” (Aberer et al. 2013) or which sit on extremely long terminal branches (Mai and Mirarab 2018). Similarly, phylogenomic studies of the relationships at a specific branch of the tree can benefit from identifying genes with a highly decisive signal (Fong et al. 2012) or those with the signal of a long branch separating the taxa in question (Chen et al. 2015). Given that multiple factors can affect branch-length estimates, using a mixture of methods that identify possibly misleading genes as well as lineages is likely to be effective for data filtering in phylogenomics.

We found that branch support strongly explains our measures of topological accuracy. Previous work has shown that gene trees with high bootstrap branch supports are associated with greater nodal support values in species-tree inferences (Blom et al. 2016). The branch-support metric used in our analyses, SH-aLRT support (Anisimova and Gascuel 2006), reflects the consistency in the signal of a given branch across the sites in the data set. High values indicate that there is a concordant signal across a large number of the informative sites. Low values can occur in genes that have few informative sites, have high degrees of rate heterogeneity across sites, or that are affected by saturation or intragenic recombination. Therefore, mean branch support is likely to provide another useful diagnostic of phylogenetic accuracy across genes. However, the relative performance of different branch-support metrics in indicating phylogenetic accuracy is yet to be explored (e.g., Lemoine et al. 2018; Minh et al. 2018).

The results of our study offer a basis for developing a framework for phylogenomics that prioritizes the inclusion of genes with a signal of limited variation in root-to-tip distances and a signal of topology that is highly concordant across sites. Our results suggest that the overall substitution rate is of limited importance as long as the evolutionary process has been homogeneous across lineages from the root of the process to the present. Potential avenues for future research include exploring the accuracy in the signal of particular types of deviation from a constant evolutionary rate across lineages, exploring the importance of model adequacy when estimating branch lengths, or comparing the performance of various metrics of branch support for predicting phylogenetic accuracy. Further examination of the correlates of reliable phylogenetic signal will be useful for selecting genes for phylogenomic analyses.

## Acknowledgements

This work was supported by funding from the Australian Research Council to S.Y.W.H. (grant FT160100167) and to D.A.D. (grant DE190100544).

---

**Supplementary Figure S1.**
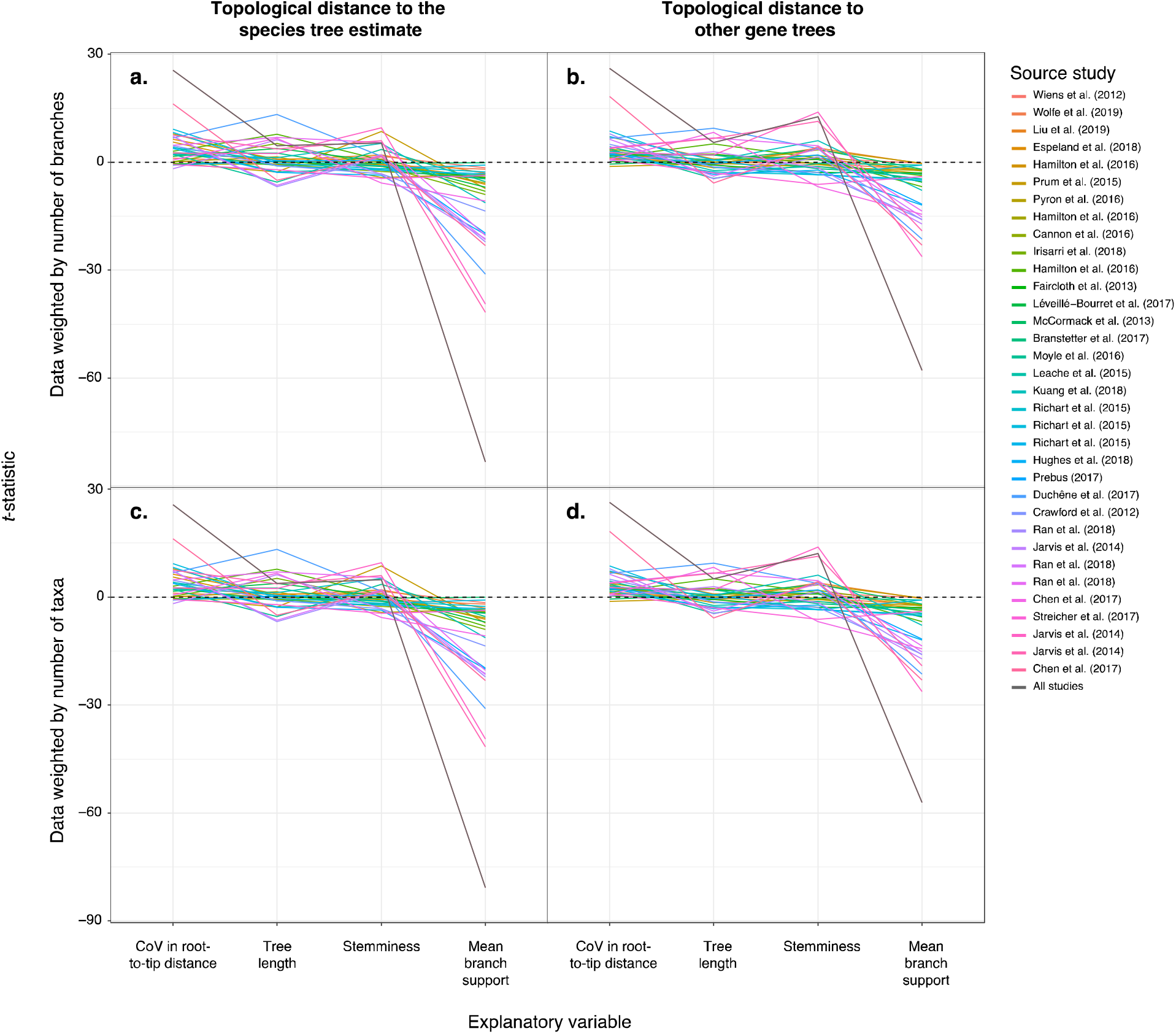
Summary *t*-statistic for multiple regression tests of the association between five explanatory variables describing branch lengths and each of two response variables: (a, c) topological distance between gene trees and the inferred species tree; and (b, d) mean distance from each gene tree to all other gene trees. Rows of panels indicate the results of analyses where regression samples (genes) were weighted by the number of branches (a, b) and number of taxa (c, d) in respective studies. The legend lists the studies in ascending order of number of genes in the data set (see Table 1 for details).

**Supplementary Figure S2.**
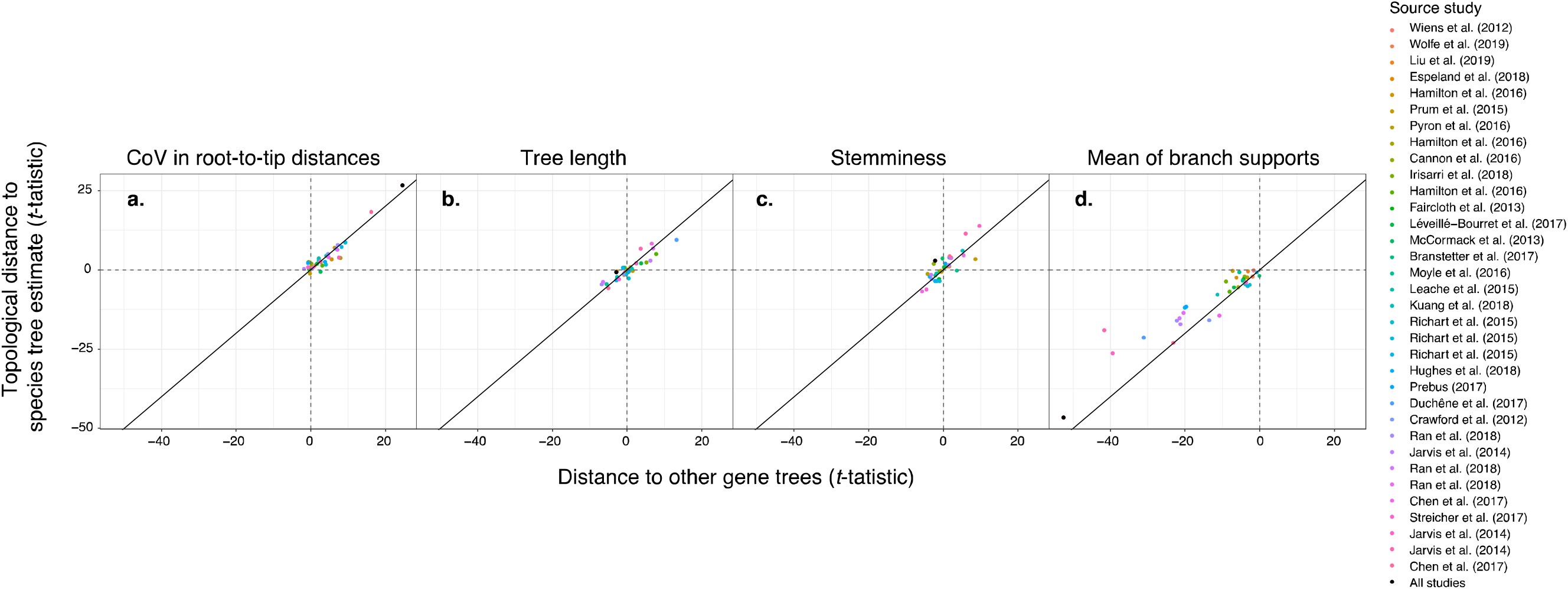
Relationship between results of multiple regression models in which the response variable was the distance to the estimated species tree (y-axis) and the mean distance to other gene trees (x-axis). Panels (a-e) show the association for each of the five explanatory regression terms included. The black point indicates the results of the regression model that included the complete data set with the source study of each genes included as a random factor. Studies in the legend are shown in ascending order of number of genes included.

